# Taxonomic Bias and Traits of the Global Amphibian Pet-Trade

**DOI:** 10.1101/645259

**Authors:** Nitya Prakash Mohanty, John Measey

**Author notes:** Corresponding Author: Nitya Prakash Mohanty.

## Abstract

The pet-trade is recognized as the major pathway for amphibian introductions worldwide, yet our understanding of the trade is limited. In this study, we systematically assess amphibian species in the pet-trade, i) characterising taxonomic bias and ii) evaluating species-traits as predictors of traded species and trade volume. We collated a global list of 443 traded amphibians and a regional dataset on trade volume. Species-traits (body size, native range size, clutch size, and breeding type) and conservation status, were considered as predictors of traded species and volume. We found a strong bias for certain Families, along with a preference for large-bodied and widely distributed species with a larval phase, in the pet-trade. However, species-traits performed poorly in predicting trade volume of pet amphibians in the USA. The identified species-traits and taxonomic bias of the trade is used to predict species likely to be traded as pets in the future.

## Introduction

Trade in live amphibians has increased drastically over the last few decades [1]; trade for pets in particular is responsible for the majority of amphibian introductions beyond their native range [2]. Van Wilgen and colleagues [3] recently recorded 263 species of amphibians with extra-limital populations, including those in trade, captivity or with non-established populations. Current patterns of invasions are driven by historic introductions (‘invasion debt’; [4]), and thus current trade will likely influence future invasions. Current lists of extralimital species [3] suggest that future invasions will encompass a broader taxonomic diversity than is currently known [5]. Therefore, it is essential to understand characteristics of species in the pet-trade and its taxonomic biases.

The amphibian pet-trade has recently received attention from studies aiming to, characterise trade regionally [6,7] and internationally [8,9,10], and to inform risk assessments [11]. However, the amphibian pet-trade still remains poorly understood. Regional trade data is available only for a limited set of countries [8,12]. Geographically, pet ownership and trade in Asia is understudied [13,14]. Research is also taxonomically biased, with trade in species listed by the Convention on International Trade in Endangered Species of Wild Fauna and Flora (CITES) being relatively well documented, even though other species may account for a much larger component of international trade [1].

Given that trade is dynamic and new species enter the trade frequently [1,6], it is essential to move beyond currently traded species and understand which species are likely to be traded in the future. Broad-scale predictors of traded species, such as life-history traits have been used to understand pet-trade dynamics (e.g. reptiles [15]; birds [16,17]). Species-traits associated with characteristics of the amphibian pet-trade (species traded and trade volume) have not yet been assessed, although release of amphibian pets is known to be influenced by life-history traits and economic parameters [18].

The amphibian pet-trade has emerged as a subject of conservation importance from the viewpoint of invasions, overexploitation, and diseases [10]. Yet, systematic assessments of the pet-trade seldom test its predictors (but see [18]). In this study, we aim to characterize amphibian species in the pet-trade. Specifically, i) we characterise taxonomic bias in traded species and ii) evaluate life-history traits as predictors of traded species and trade volume.

## Methods

We collated a list of traded amphibians based on a literature review (Supplementary Information 1) and supplemented it with the United States Fish and Wildlife Service’s Law Enforcement and Management Information System (LEMIS) data on live imported amphibians into the USA from 2013 to 2018. Based on the ‘purpose’ variable associated with the imports, we considered only ‘commercial’, ‘breeding in captivity’ and ‘personal’ categories as those pertaining to the pet-trade. Since our interest was in species-level traits as predictors, only import records with identified species were retained, discarding taxa identified up to genus-level and unspecific categorization such as ‘non-CITES’. We corrected for taxonomy based on Frost [19] and assigned an Order, Superfamily, and Family to each species. Volume of trade is rarely recorded and LEMIS data for the USA provides a unique opportunity to make further inferences about trade (e.g. [12]), especially on species’ popularity in the pet-trade. The total number of individuals imported for each species into the USA from 2013 to 2018 served as a measure of ‘trade volume’.

Species-traits for amphibians were collated mainly from the AmphiBIO database [20]. This was further supplemented by data from Allen et al. [21] and AmphibiaWeb (https://amphibiaweb.org/). We selected traits with data available for a majority of species, which were likely to influence pet ownership and trade. Range size may influence availability for trade [22] and therefore data on native range and global range (i.e. including non-native range) was obtained by geoprocessing polygons from the IUCN spatial database [23]. Body size is likely to determine the suitability of species for the pet-trade [24]; extremely small or large body sizes may be avoided due to reduced detection and increased costs of housing, respectively [18]. Clutch size and breeding mode (direct developing, larval, or viviparous) have direct bearing on the ease of captive breeding.

Trade status of species, as recorded by CITES, and conservation status according to the International Union for Conservation of Nature’s (IUCN) Red List were also considered, as these variables could influence trade volume [17]. Each species was assigned a CITES Appendix (I, II, III) or recorded as ‘non-listed’; following Vall-llosera & Cassey [17], IUCN ‘Near-threatened’ and ‘Least Concern’ species were categorized as ‘non-threatened’, and the rest as ‘threatened’, while treating ‘Data Deficient’ as a category of its own. Data on all chosen predictors were available for a sizable number of amphibians for trade status (n = 1388) and trade volume analyses (n = 173). We did not include some potentially relevant traits (climate, diet, longevity, offspring size) as too few species are scored in the AmphiBIO database.

### Data Analyses

To evaluate taxonomic bias in representation of amphibian Orders, Superfamilies or Families in traded species, number of species at each taxonomic level was compared with the total number of known amphibian species [19], with respect to a random expectation generated using the hypergeometric distribution (see [3]). Taxa outside the 95% confidence intervals were deemed either over- or under-represented in our sample of traded species. To assess the predictors of traded species and trade volume, we constructed a mixed-effects logistic regression model in the lme4 package [25]. Prior to the analyses, data on both body size, clutch size, and range sizes (native and global range) were log-transformed. We controlled for potential taxonomic dependence by using ‘Family’ as a random effect in our analysis on traded species and ‘Superfamily’ for trade volume. We did not include ‘CITES’ status as a predictor of traded species, but only for trade volume analyses. Model selection was based on the Akaike Information Criterion (AIC), supplemented by a measure of model fit (R^2^_GLMM_; [26]). Based on the parameter estimates of the best model, ‘trade scores’ (likelihood to be traded) were computed for all species and scaled for comparison. All statistical analyses were carried out in R (version 3.5.3; [27]), whereas, spatial computations were executed in ArcMap (version 10.6.1; [28])

## Results

### i) Taxonomic bias

Our literature review and US import records resulted in a total of 443 uniquely identifiable species in the amphibian pet-trade from 1971 to 2018 (Supplementary Information 1). Frogs (Order Anura) constituted the majority of the traded species (n = 262), followed by salamanders (Caudata = 47) and caecilians (Gymnophiona = 3). An analysis accounting for all extant taxa revealed a taxonomic bias in traded amphibians, with over- and underrepresented taxa at Order, Superfamily, and Family level (Fig. 1). At the Order level, Caudata were over-represented in traded taxa, whereas Anuran and Gymnophiona were under-represented (Fig. 1a). Traded species were also biased towards the Superfamilies Salamandoidea, Pipoidea, Dendrobatoidea, and Discoglossoidea (Fig. 1b). The Families Dendrobatidae, Mantellidae, Hyperoliidae, Pipidae, Ambystomatidae, and Salamandridae contributed disproportionately high species to the trade (Fig. 1c).

**Figure 1.**
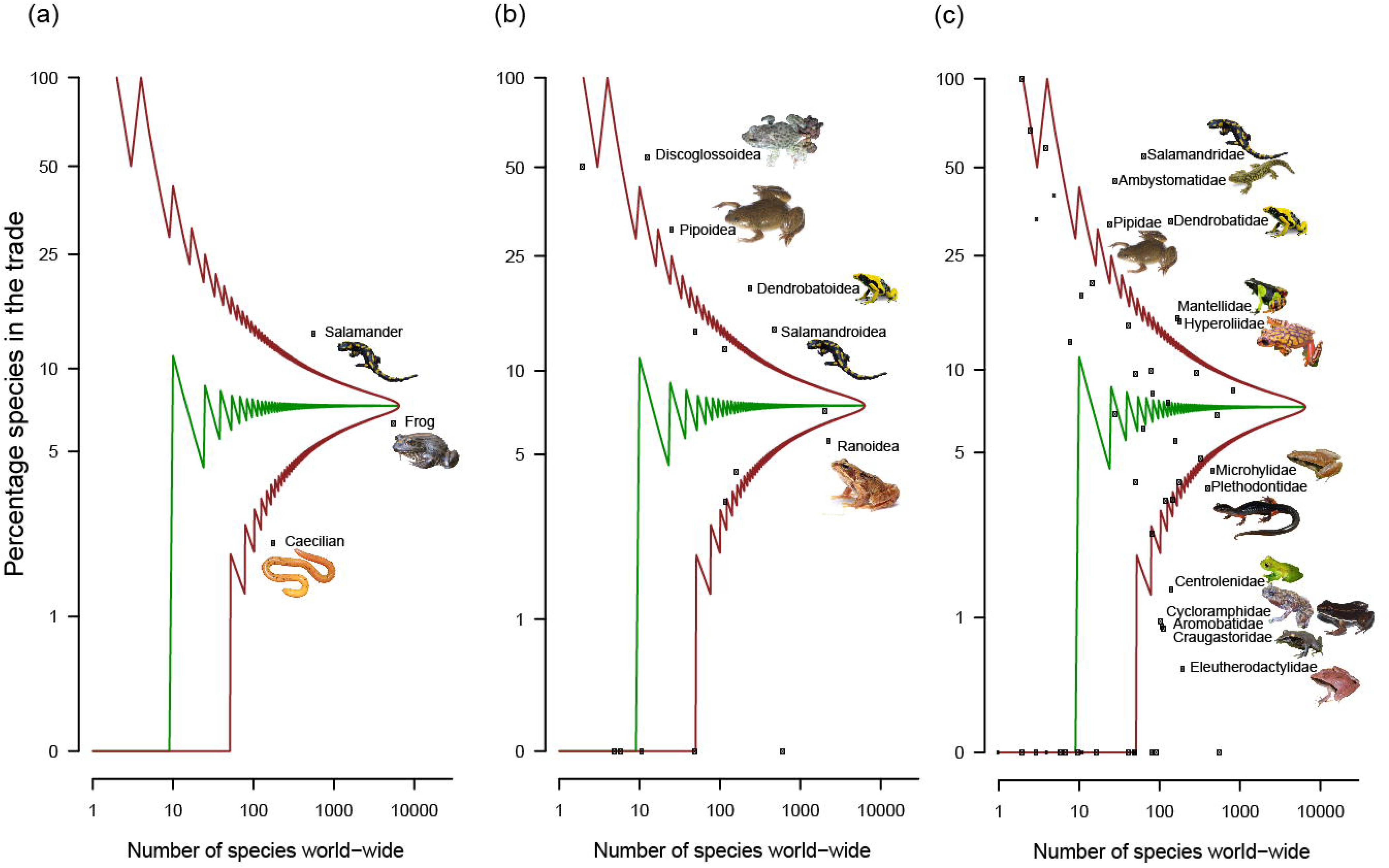
Taxonomic patterns in amphibian (A) Orders, (B) Superfamilies, and (C) Families present in the global amphibian pet-trade. The median (green line) and 95% confidence intervals (brown lines above and below), adjusted for multiple comparisons, were estimated from the hypergeometric distribution. The points that fall between the 95% confidence intervals are not significantly over or under-represented, relative to the number of amphibian species worldwide. Those labelled taxa that fall above the 95% confidence intervals are over-represented and those below are under-represented in our sample of traded amphibians.

### ii) Predictors of traded species and trade volume

Of the 15 candidate models built to predict traded species, the global model performed the best (Table 1). However, only body size, range size, and breeding type had significant effects, with the likelihood for species to be traded being positively influenced by body size (*β* = 1.06, SE = 0.19, *p* < 0.001), range size (*β* = 0.38, SE = 0.05, *p* < 0.001), and a ‘larval’ breeding type (*β* = 1.34, SE = 0.50, *p* = 0.007). The best model explained 59% of the variation overall (including the random effect of Family); on their own the predicting traits explained 41% of the variation (Table 1). Native range size and global range size did not differ in their explanatory ability. Based on their ‘trade scores’ (output of the best model), we produced a list of 20 species likely to be traded in the future (Table 2).

**Table 1.** Generalized linear mixed-effects models predicting amphibian species in the pet-trade with predictors body size, native range size (km^2^), breeding mode (LA – larval, VV – viviparous, DD – direct developing), and clutch size. Models run with ‘Family’ as a random effect. ΔAIC is the difference in Akaike information criterion values (AIC) between the current model and the best and weight (Akaike weight) is the relative support a model has from the data compared to the other models in the set. Marginal (*R*^2^_*m*_) and conditional (*R*^2^_*c*_) R^2^_GLMM_ are reported for each model and provide an estimate of the explained variance.

| Model | $\Delta AIC$ | Weight | $R^2_m$ | $R^2_c$ |
| --- | --- | --- | --- | --- |
| Size + Range + Breeding + Clutch + IUCN | 0.00 | 0.83 | 0.41 | 0.59 |
| Size + Range + Breeding | 3.61 | 0.14 | 0.41 | 0.59 |
| Size + Range + Clutch | 7.13 | 0.02 | 0.36 | 0.57 |
| Size + Range + IUCN | 10.00 | 0.01 | 0.34 | 0.57 |
| Size + Range | 12.83 | 0.00 | 0.34 | 0.57 |
| Range + Breeding | 54.54 | 0.00 | 0.36 | 0.51 |
| Range + IUCN | 59.52 | 0.00 | 0.28 | 0.48 |
| Range | 66.62 | 0.00 | 0.28 | 0.47 |
| Size + Breeding | 106.69 | 0.00 | 0.26 | 0.51 |
| Breeding + Clutch | 127.67 | 0.00 | 0.25 | 0.46 |
| Size | 127.67 | 0.00 | 0.19 | 0.51 |
| Clutch | 135.05 | 0.00 | 0.17 | 0.42 |
| Breeding | 194.60 | 0.00 | 0.13 | 0.32 |
| IUCN | 197.97 | 0.00 | 0.06 | 0.29 |
| Null | 226.99 | 0.00 | 0.00 | 0.27 |

**Table 2.**
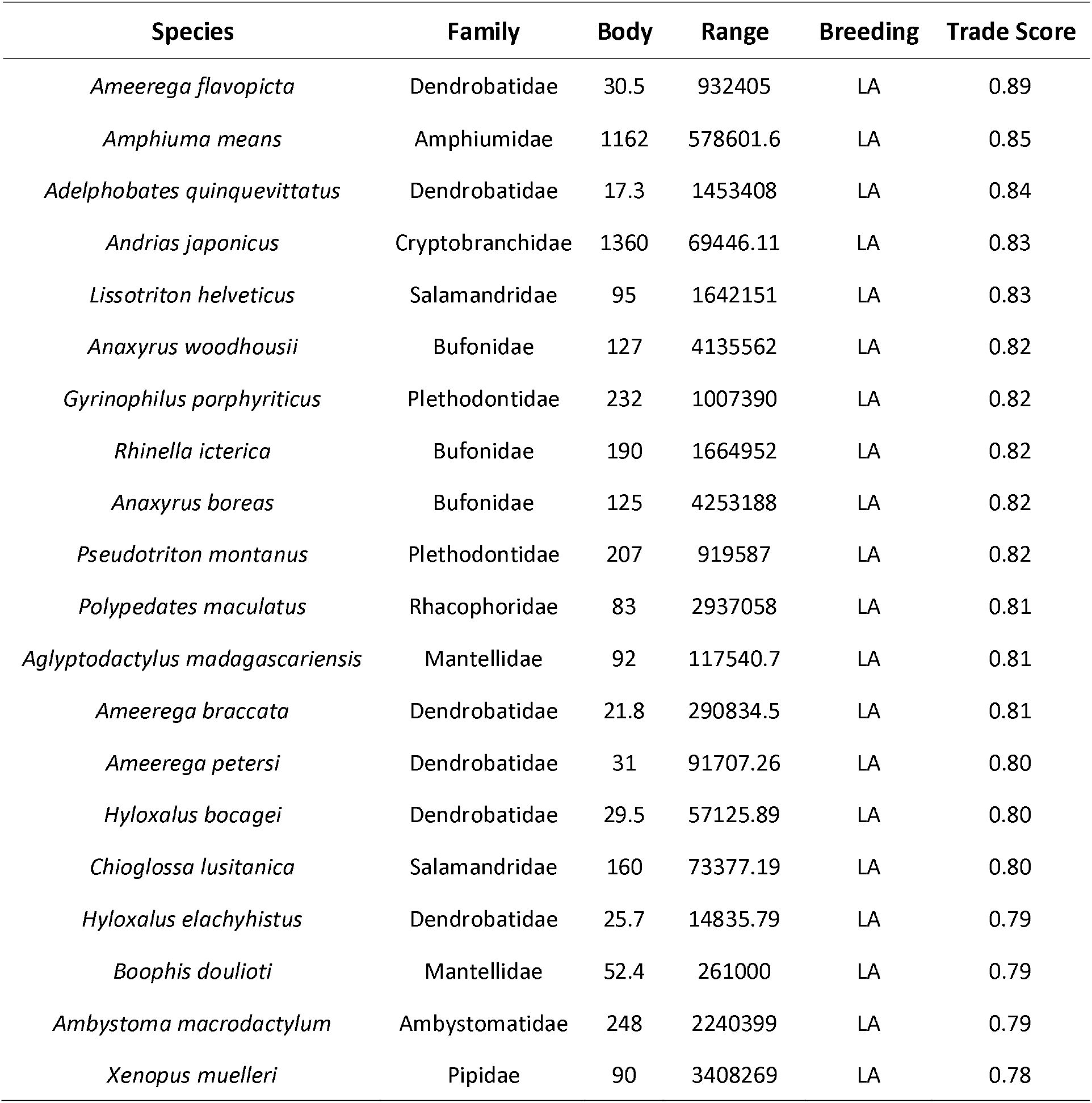
List of example amphibian species that are likely to be traded as pets in the future, based on parameter estimates (‘trade score’) of the selected species-trait based model, with associated body size (mm), native range size (km^2^), and breeding mode (LA – larval). ‘Trade score’ is standardized (from 0 to 1) to enable comparisons of candidate species for the pet-trade.

| Species | Family | Body | Range | Breeding | Trade Score |
| --- | --- | --- | --- | --- | --- |
| *Ameerega flavopicta* | Dendrobatidae | 30.5 | 932405 | LA | 0.89 |
| *Amphiuma means* | Amphiumidae | 1162 | 578601.6 | LA | 0.85 |
| *Adelphobates quinquevittatus* | Dendrobatidae | 17.3 | 1453408 | LA | 0.84 |
| *Andrias japonicus* | Cryptobranchidae | 1360 | 69446.11 | LA | 0.83 |
| *Lissotriton helveticus* | Salamandridae | 95 | 1642151 | LA | 0.83 |
| *Anaxyrus woodhousii* | Bufonidae | 127 | 4135562 | LA | 0.82 |
| *Gyrinophilus porphyriticus* | Plethodontidae | 232 | 1007390 | LA | 0.82 |
| *Rhinella icterica* | Bufonidae | 190 | 1664952 | LA | 0.82 |
| *Anaxyrus boreas* | Bufonidae | 125 | 4253188 | LA | 0.82 |
| *Pseudotriton montanus* | Plethodontidae | 207 | 919587 | LA | 0.82 |
| *Polypedates maculatus* | Rhacophoridae | 83 | 2937058 | LA | 0.81 |
| *Aglyptodactylus madagascariensis* | Mantellidae | 92 | 117540.7 | LA | 0.81 |
| *Ameerega braccata* | Dendrobatidae | 21.8 | 290834.5 | LA | 0.81 |
| *Ameerega petersi* | Dendrobatidae | 31 | 91707.26 | LA | 0.80 |
| *Hyloxalus bocagei* | Dendrobatidae | 29.5 | 57125.89 | LA | 0.80 |
| *Chioglossa lusitanica* | Salamandridae | 160 | 73377.19 | LA | 0.80 |
| *Hyloxalus elachyhistus* | Dendrobatidae | 25.7 | 14835.79 | LA | 0.79 |
| *Boophis doulioti* | Mantellidae | 52.4 | 261000 | LA | 0.79 |
| *Ambystoma macrodactylum* | Ambystomatidae | 248 | 2240399 | LA | 0.79 |
| *Xenopus muelleri* | Pipidae | 90 | 3408269 | LA | 0.78 |

The USA imported at least 3,655,620 live amphibians for the pet-trade, belonging to 283 species, between 2013 to 2018 (Supplementary Information 2). Three models were selected (∆AIC<2) to explain trade volume, with predictors breeding mode, CITES, and IUCN status. However, these predictors explained only 1.5% of the variation in trade volume (Supplementary Information 2).

## Discussion

In this study, we collate a comprehensive list of amphibian species in the global pet-trade. We find a strong bias for certain Families, along with a preference for large-bodied and widely distributed species with a larval phase, best characterise the global trade in amphibians. Our analyses reveal that a few amphibian Families, with high species diversity (e.g. Mantellidae, Dendrobatidae, Hyperoliidae, Salamandridae), contribute disproportionately to the trade. Species within a Family are likely to have broadly similar traits that make them candidates for the pet-trade [17]. Along with this taxonomic bias, the identified species-traits can predict new species likely to enter the trade. Our example list of species based on predictions of trait-based models (Table 2) suggests that future traded species are likely to be from already overrepresented Families (e.g. Dendrobatidae).

The modelling results indicate that traded species are associated with species-trait preference (e.g. large body size), human sampling effort (e.g. large range size), and husbandry practices (e.g. indirect development). Body size is known to be positively associated with intentional introduction of amphibians [22]. This result is unsurprising as most intentional introductions are through the pet-trade [2] and large bodied-species are more likely to be released [18]. Although large body size does not lead to higher success of establishment or spread [21], it is predictive of greater impact in invasive amphibians [29]. Tingley et al.[22] noted the influence of large range size on introduction probability and attributed it to increased opportunities for sampling by humans. Additionally, we posit that a larger range is also likely to include more countries, which may increase the chances of a species’ trade being facilitated, overcoming regulatory restrictions. Species with larval offspring are likely to be cheaper to raise as compared to direct developing species. Thereafter, most captive bred pet amphibians are imported as sub-adults (e.g. [12]). The minimal effect of conservation status (‘IUCN’) on pet-trade of species has been previously documented with birds [17].

The species-traits considered in our analyses performed poorly in predicting trade volume of pet amphibians in the USA. Human aspects of the trade, such as availability of skills for captive breeding, exemption from regulations, and economics of scale are likely to drive trade, particularly volume (see [17] for a detailed discussion). Species attributes not considered in our analyses, such as colour, ornamentation, rarity, and perceived cost of ownership [15,30], may also account for unexplained variation in traded species and their volume. However, information on these traits are not readily available for many amphibian species. Amphibian trait data also suffers from incompleteness, limiting the number of species that can be considered [20,21].

Recent studies have attempted to understand pet ownership and stakeholder perception of pet-trade management [14,31]. Future research must systematically assess human motivations for pet ownership and release, preferences for traits to better hone predictions of which species are likely to be traded, and better inform risk assessments.

## Acknowledgements

We would like to thank the USFWS for sharing data on amphibian imports; Martin Schlaepfer, Oliver Stringham, Hollis Dahn, James Baxter-Gilbert and Carla Wagener for valuable inputs to the study.

## Author contributions

N.P.M and J.M. conceived the study. N.P.M. collated and analysed the data. N.P.M. and J.M. wrote the manuscript and approved the final version of the manuscript.

## Accessibility

All data associated with the manuscript are uploaded as part of electronic supplementary material, S3.

## Ethical statement

No ethical approval was required for this study.

